# Endocannabinoid signaling is a critical link between circadian desynchronization and metabolic dysfunction

**DOI:** 10.1101/2025.09.29.678590

**Authors:** Brennan A. Baca, Giancarlo E. Denaroso, Said Akli, Gregory L. Pearson, Jiexin Wang, Nicole P. Bowles, Derrick Phillips, Walker Sorensen, Catherine Hume, Matthew N. Hill, Ilia N. Karatsoreos

## Abstract

It is well documented that disruption of circadian rhythms can cause metabolic dysregulation, but the specific mechanisms involved remain unclear. Our findings demonstrate that the negative metabolic effects of environmental circadian desynchronization (ECD) are dependent upon the cannabinoid receptor 1 (CB1r). The endocannabinoid system has not previously been implicated in mediating the effects of circadian disruption. We showed that ECD induced a positive correlation between the levels of the endocannabinoids AEA and 2-AG in both plasma and liver. While global CB1r knockout protects against the metabolic effects of ECD, behavioral and physiological response to ECD was strikingly similar between WT and CB1r KO mice and could not account for their distinct metabolic outcomes. Using liver-specific CB1r KO mice, we further specified that the ECD-induced metabolic hormone disruption, but not weight gain, is mediated through liver CB1r signaling. Finally, we showed that ECD upregulated transcription of genes involved in oxidative phosphorylation in the liver of WT, but not liver-specific CB1r KO mice. In summary, ECD led to modular metabolic dysfunction through CB1r signaling in multiple tissues, with the liver playing a critical role.

## Introduction

Circadian rhythms are near-24hr rhythms present in nearly all tissues and physiological processes^1^, including cellular and organismal metabolism. These rhythms play an important role in anticipating recurring changes in the environment, thus enabling cells, organs, and whole organisms to tune physiology and behavior to predicted changes in physiological demands. Under “normal” conditions, circadian rhythms are adaptive. However, in contexts where light/dark cycles are misaligned with the endogenous clock, such as shift work or jet lag, endogenous circadian rhythms become desynchronized from rhythmic environmental cues. This has been implicated in poor health outcomes, including metabolic disorder^2,3^. Despite the widespread prevalence of circadian rhythm disruption and metabolic disorder in modern society^4^, the potential mechanisms that link them remain poorly understood^5^.

There are myriad signaling pathways that could link circadian rhythms and metabolism. Recent research provides compelling evidence suggesting the endocannabinoid system may interface with both circadian rhythms and metabolic function. The endocannabinoid system is canonically comprised of two ligands – anandamide (AEA) and 2-arachidonylglycerol (2-AG) – as well as two primary receptors – cannabinoid receptor 1 (CB1r) and 2 (CB2r) – that, together, regulate functions spanning neurotransmission, immunity, and metabolism^6,7^. CB1r is expressed nearly ubiquitously in mammalian tissues and is known to act both centrally and peripherally to promote food seeking and efficient energy storage (e.g., adipogenesis)^8,9^. In addition, high-fat diet leads to a rapid increase in endocannabinoids^10^ and obesity through a CB1r-dependent mechanism^11^. Intriguingly, people with obesity exhibit both elevated levels and disrupted daily rhythms of circulating endocannabinoids^12–15^. The liver is uniquely situated at the intersection of these systems – it regulates metabolic function, it is endocannabinoid competent^16,17^, and is the organ with perhaps the highest proportion of rhythmically expressed gene transcripts^18^.

We have previously demonstrated that environmental circadian desynchronization (ECD) via exposure to 10 weeks of housing in a 20hr day (10hr light, 10hr darkness) leads to weight gain, elevated plasma leptin and insulin, as well as a variety of neurocognitive effects^19^, with sleep and immune effects visible after just 4 weeks^20^. In the present study, we used a 5-week exposure to ECD to study how early behavioral and physiological changes may contribute to subsequent metabolic disruptions. We first show that ECD induced a significant and positive correlation between endocannabinoids AEA and 2-AG in both plasma and liver tissue, uncovering a potential link between ECD, endocannabinoids, and metabolic disruption. To understand if CB1r signaling is necessary for ECD-induced metabolic dysfunction, we undertook continuous, longitudinal analysis of the behavior and physiology of both wildtype (WT) and global CB1r knockout (KO) mice in Control and ECD conditions. We uncovered that global CB1r signaling was necessary for ECD-induced weight gain and metabolic hormone dysregulation. Surprisingly, this protective effect of CB1r KO appeared to be independent of changes to gross or rhythmic behavior and metabolic physiology, as many of these measures were similarly affected by ECD in both WT and CB1r KO mice. Intriguingly, and consistent with the metabolic role of the liver, our data point to hepatocyte CB1r as a key site of ECD-induced metabolic disruption. Finally, our analyses suggest that the ECD-induced metabolic phenotype may result from altered hepatic mitochondrial function. Collectively, our findings establish a clear role of the endocannabinoid signaling system in the development of ECD-induced metabolic disruption.

## Results

### ECD alters endocannabinoid coregulation and induces a CB1r-dependent metabolic phenotype

We assessed the impact of ECD on endocannabinoids by measuring levels of AEA and 2-AG in both plasma and liver of WT mice in Control and ECD conditions. We found several main effects, including an effect of time with respect to the light cycle (*zeitgeber* time, ZT) for plasma 2-AG and liver AEA in males, and a slight but significant effect of ECD in female plasma AEA (Figures S1A-S1H).

While ECD did not cause a marked change in overall endocannabinoid levels, linear regression analysis revealed that ECD induced a positive, within-animal correlation between AEA and 2-AG levels in both the plasma and liver of male mice (Figures 1A and 1B). We observed a similar effect in female plasma, and a trend in female liver tissue that did not reach statistical significance (Figures S1I and S1J). Thus, ECD induced a positive correlation between AEA and 2-AG in plasma and liver tissue, while having little effect on their absolute concentrations.

**Figure 1:**
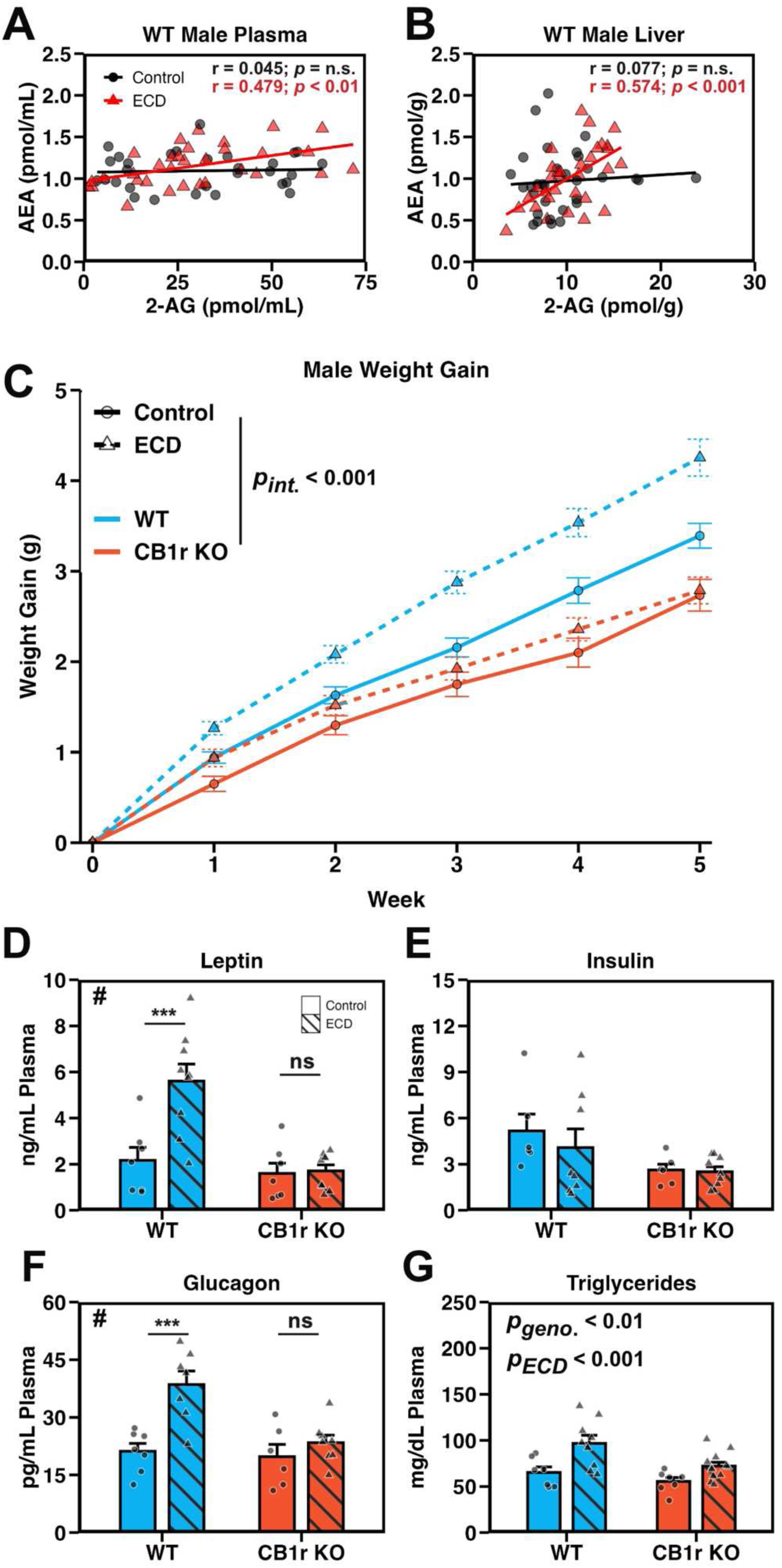
ECD alters endocannabinoid coregulation and induces a CB1r-dependent metabolic phenotype. **A and B:** Scatterplot of AEA and 2-AG levels in plasma (A) and liver (B) from male mice under Control (black circles) and ECD (red triangles) conditions and their linear regressions (lines). Samples were collected at various time points (one of: ZT 0, 4, 8, 12, 16, or 20) and collapsed for correlation analysis. *n* = 30-35/experimental condition/tissue. **C:** Weight gain of male mice recorded manually every 7 days. Data were analyzed using a linear mixed effects model (*p*_Genotype x Condition x Week_ < 0.001) and within-group follow-up analysis of the effect of ECD performed using estimate of marginal means (*p*_WT_ < 0.001; *p*_KO_ = 0.205). *n* = 54-71/genotype/experimental condition/week. **D – G:** Plasma levels of leptin (D; *p*_Int_ = 0.005, *p*_WT_ < 0.001, *p*_KO_ = 0.887), insulin (E; *p*_Int_ = 0.396), glucagon (F; *p*_Int_ = 0.018, *p*_WT_ < 0.001, *p*_KO_ = 0.352) and triglycerides (G; *p*_Int_ = 0.241) from male mice collected at ZT 12. Significant interaction between genotype and ECD indicated by “#”, significant post-hoc t-test indicated by “*”. *n* = 6-12/genotype/experimental condition. **Stats:** Data are reported as mean +/- the standard error of the mean. For endocannabinoid correlations (A and B), a linear regression was fit using the lm() function in R, and the resultant *p*-value and Pearson’s correlation coefficient (r) value were reported. For weight gain (C), main effects and follow up comparisons were performed using a mixed effects model and estimate of marginal means, respectively (see methods). For plasma metabolite assessment (D-G), we performed a two-way ANOVA (or non-parametric ART ANOVA if assumptions were not met; see methods). Following a significant interaction term, follow-up post-hoc t-tests were performed and *p*-values adjusted using the Bonferroni method. \**p* < 0.05, \*\**p* < 0.01, \*\*\**p* < 0.001

We then tested whether signaling through CB1r contributed to metabolic disruption following ECD by exposing male and female global CB1r WT and KO mice to ECD. There was a significant three-way interaction between experimental condition, genotype, and week of experiment on weight gain in male mice (Figure 1C). Follow-up, pairwise comparisons of estimated marginal means revealed that WT, but not CB1r KO males, gained excessive weight following ECD. ECD did not induce excessive weight gain for females of either genotype (Figure S2A). To better understand how this weight gain phenotype and the CB1r KO-dependent protection occurred, we focused subsequent experiments on male mice.

Using a panel of assays, we found that ECD elevated plasma levels of leptin and glucagon in WT, but not KO males, and did not alter insulin levels in either genotype (Figures 1D-1F). Plasma triglycerides were elevated by ECD and reduced by CB1r KO (Figure 1G). Gastric inhibitory peptide (GIP) was elevated by ECD in both WT and CB1r KO males, while amylin and peptide YY (PYY) were unchanged by ECD (Figure S3A). ECD elevated inflammatory markers monocyte chemoattractant protein 1 (MCP1) and interleukin-6 (IL-6) in WT, but not CB1r KO (Figure S3A). There was a trend toward reduced tumor necrosis factor alpha (TNF-α) in WTs, and an increase in CB1r KOs, following ECD (Figure S3A). Liver triglyceride levels, which are typically elevated following significant metabolic disruption and indicate hepatic steatosis^21^, were unchanged by ECD in both genotypes (Figure S3C).

Together, these data suggest that a 5wk exposure to ECD induced an early metabolic phenotype in WT mice, and that signaling through CB1r was necessary for the development of this phenotype.

### ECD affects the temporal organization of physiological processes independent of CB1r signaling

To understand how ECD alters energy balance, and elucidate any role for CB1r in this process, we performed continuous home-cage phenotyping during the 5wk exposure to Control or ECD conditions. This allowed us to monitor feeding, drinking, and locomotion as well as gas exchange, permitting the calculation of energy expenditure (Kcal/hr) and respiratory exchange ratio (RER), a unitless value indicating composition of whole-body macronutrient substrate oxidation.

Even though ECD led to weight gain in WT males (Figure 1), we found that it had no significant effect on gross energy intake or energy expenditure in either genotype (Figures 2A and 2B). While CB1r KO males ate less than their WT littermates, this effect was absent after correcting for body weight (Figure S4) and could not explain why ECD caused weight gain in WT, but not CB1r KO, males.

**Figure 2:**
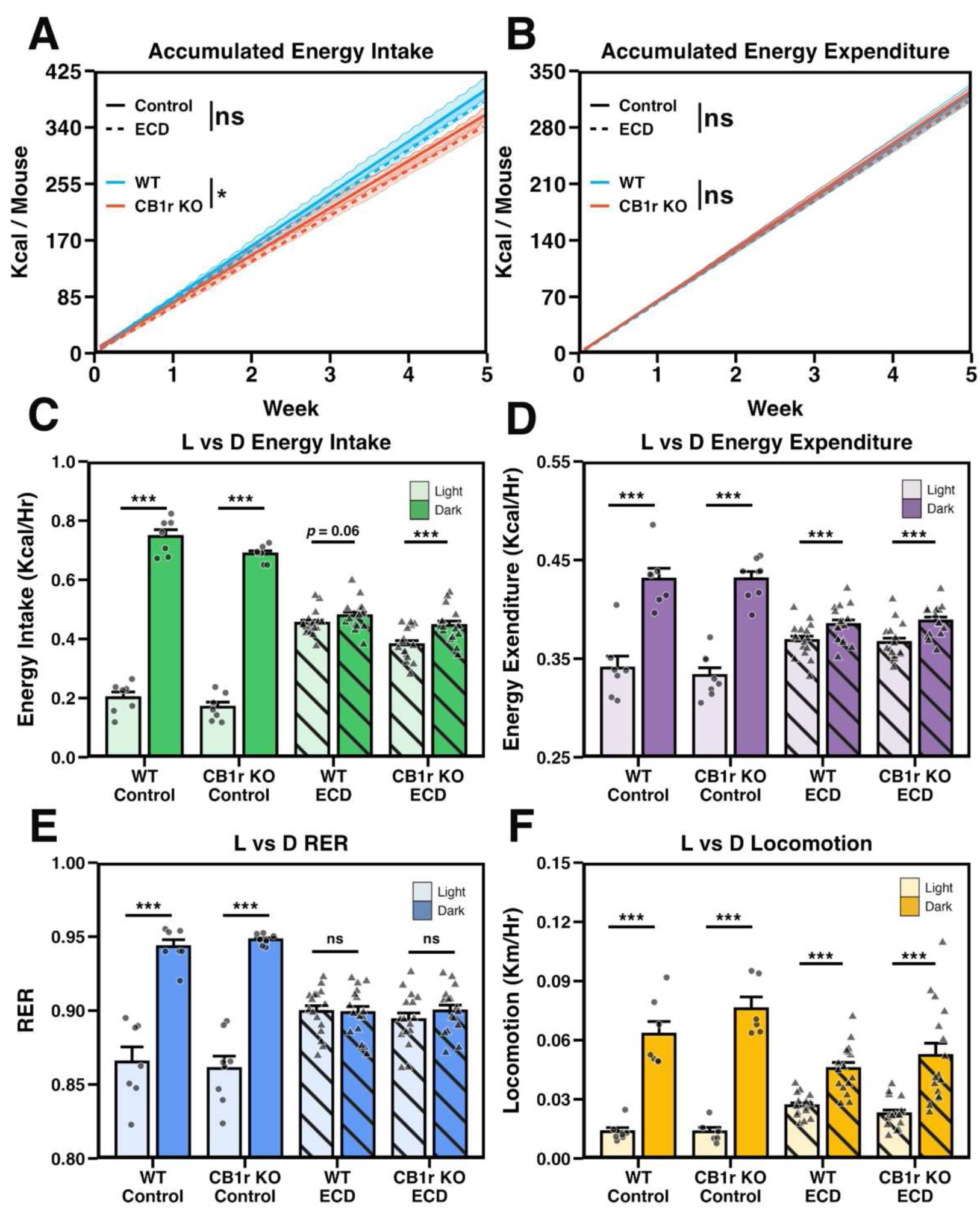
ECD affects the temporal organization, but not gross organization, of physiological processes independent of CB1r signaling. **A and B:** Cumulative energy intake (A; *p*_Genotype x Condition x Week_ = 0.868) and energy expenditure (B; *p*_Genotype x Condition x Week_ = 0.948) in Control and ECD conditions. Data are shown as mean (solid or dashed line) +/- the standard error of the mean (shaded region). Results of interaction term between genotype and week or experimental condition and week are shown within figure by “*”. *n*_Control_ = 7/genotype, *n*_ECD_ = 16/genotype. **C – F:** Average energy intake (C; *p*_Genotype x Condition x Phase_ = 0.0556), energy expenditure (D; *p*_Genotype x Condition x Phase_ = 0.778), respiratory exchange ratio (“RER”; E; *p*_Genotype x Condition x Phase_ = 0.829) and locomotor activity (F; *p*_Genotype x Condition x Phase_ = 0.446) organized by light phase in Control and ECD conditions. *n*_Control_ = 6-8/genotype/phase, *n*_ECD_ = 16/genotype/phase. **Stats:** For cumulative daily energy intake and expenditure (A and B, respectively), main effects and follow-up comparisons were performed using a mixed effects model and estimate of marginal means, respectively (see methods). A 3-way ANOVA was used to test the effects of phase, genotype, experimental condition and their interactions, on physiological data (C-F), with planned t-tests performed to assess the difference between light and dark phase activity within each group. *p*-values were adjusted using Bonferroni method. \**p* < 0.05, \*\**p* < 0.01, \*\*\**p* < 0.001.

We then assessed the daily organization of biological processes to test for group differences in the ability to adapt to the 20hr day. As expected, feeding, locomotion, RER, and energy expenditure were greater during the dark (active) phase compared to the light phase in Control conditions (Figures 2C-2F). ECD dramatically altered the light/dark organization for all of these measures. Importantly, WT and CB1r KO males exhibited a similar response to ECD for all of these measures, as indicated by the lack of significant three-way interactions between genotype, experimental condition, and phase.

To further assess how WT and CB1r KO organized their behavior in ECD conditions, we calculated the ZT of daily maxima for locomotion, feeding, RER, and energy expenditure (Figures 3A-3H; corresponding actograms provided in Figure S5). When analyzed across all animals using the Rayleigh test of uniformity for circular data, both the magnitude (Z) and frequency of significant clustering were reduced under ECD conditions for locomotion, feeding, RER, and energy expenditure in both genotypes (Figures 3I-3L). Importantly, there were no significant effects of genotype or interactions on these measures, suggesting that WT and CB1r KO animals respond similarly to ECD.

**Figure 3:**
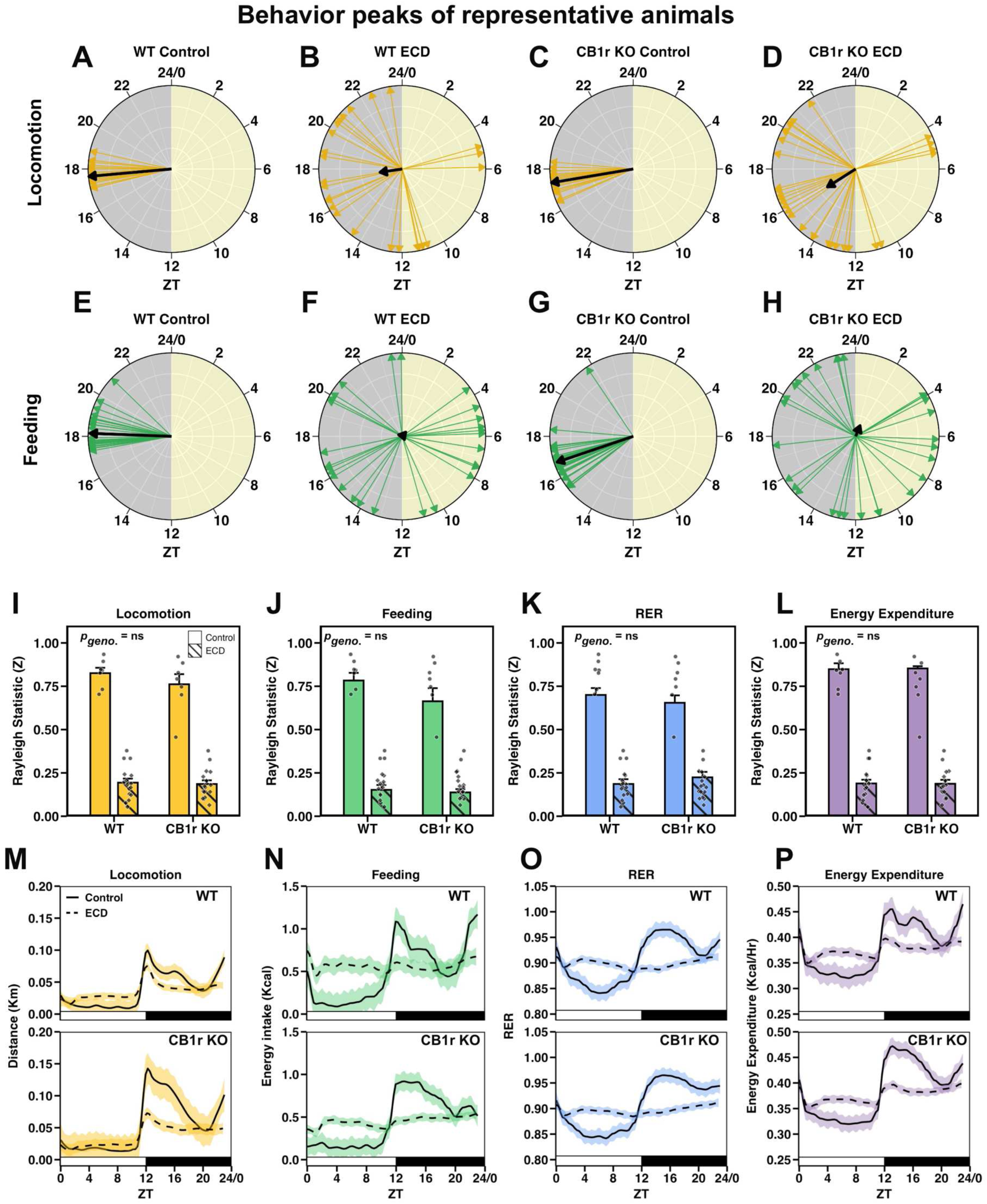
ECD misaligns behavioral and environmental rhythms independent of CB1r signaling. **A – H:** Radial plots showing *zeitgeber* time (ZT) of peak in daily locomotion (A-D) and feeding (E-H) for representative animals. Daily behavior peaks are indicated with colored arrows, and the resultant average vector is shown by a black arrow. The magnitude of the average vector is proportional to the Rayleigh statistic “Z”. **I – L:** Quantification of the Rayleigh statistic (Z) computed for the daily peak timing of locomotion (I; *p*_Int_ = 0.389), feeding (J; *p*_Int_ = 0.177), respiratory exchange ratio (K; *p*_Int_ = 0.265) and energy expenditure (L; *p*_Int_ = 0.751) of all animals in Control and ECD conditions. Significant results of the Rayleigh test of uniformity for circular data are shown as circles while non-significant results are shown as diamonds. Under Control conditions for both WT and CB1r KO groups, all animals (7/7 per group) exhibited significant clustering for each behavior. In ECD conditions, 3/16 WTs and 2/16 CB1r KOs exhibited significant clustering for each of the four measures. *n*_Control_ = 7/genotype/measure, *n*_ECD_ = 16/genotype/measure. **M – P:** Average locomotion (M), feeding (N), RER (O), and energy expenditure (P) of all WT and KO males (top and bottom panels, respectively). Data over the 5wk experiment were first averaged within animal and ZT, and then the mean of all animals (solid line) and standard error across animals (shaded region) were computed. Solid lines indicate Control and dashed lines indicate ECD. *n*_Control_ = 7/genotype, *n*_ECD_ = 16/genotype. **Stats:** A 2-way ANOVA was used test the effects of genotype, experimental condition, and their interactions on the Rayleigh statistic (Z).

Finally, we visualized the average locomotion, feeding, RER, and energy expenditure of all male mice with respect to the light/dark cycle (Figures 3M-3P). Under Control conditions, all four of these measures exhibited reliable peaks shortly after lights off and reached their nadir during the light phase, as expected. This was severely blunted in ECD for both WT and CB1r KO males. In summary, the effects of ECD on gross metabolic physiology were largely equivalent across WT and CB1r KO animals.

### ECD reduces the amplitude and precision of physiological rhythms independent of CB1r signaling

To test for group differences in the ability to preserve endogenous near-24hr biological rhythms under ECD conditions, we performed chi-square periodogram analysis (Figures 4A-4D). To assess periodicity of locomotion, feeding, RER, and energy expenditure across all animals, we considered the two largest peaks (i.e., the primary and secondary peaks, as defined by power) that crossed the significance threshold and plotted their periods (Figures 4E-4H). We found that while all of these physiological measures exhibited precise 24hr rhythms in Control conditions, these near-24hr rhythms became less precise and exhibited less power in ECD for both genotypes. A subset of ECD animals (WT: 6/16; KO: 7/16) exhibited primary near-20hr rhythms in locomotion, which was likely a result of masking, in which an animal’s behavior is directly altered by, but not entrained to, external cues. Notably, there were no observable group differences in the response of behavioral rhythms to ECD between WT and CB1r KO mice, eliminating this as a likely source of CB1r KO-mediated metabolic protection.

**Figure 4:**
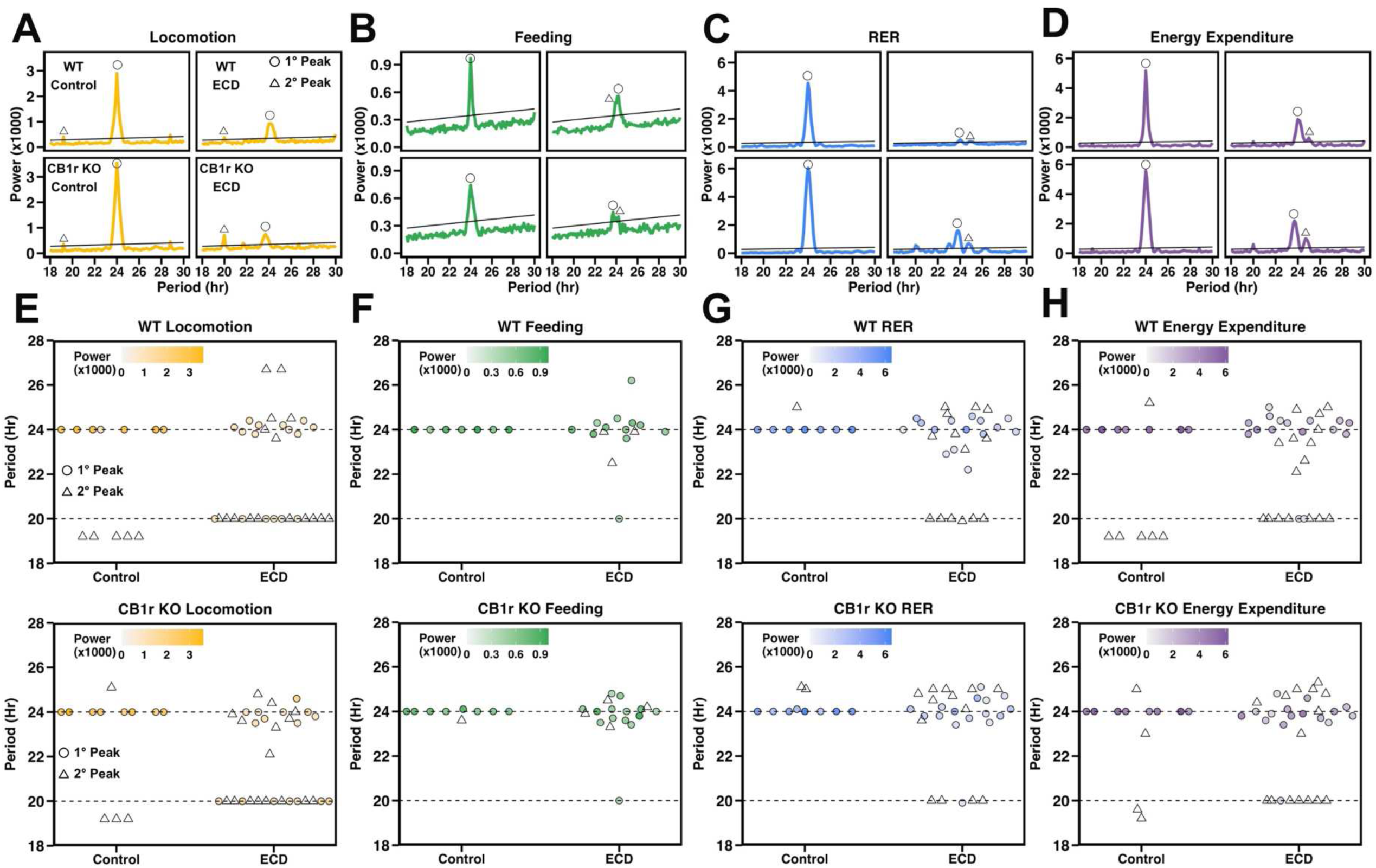
ECD reduces the amplitude and precision of physiological rhythms independent of CB1r signaling. **A – D:** Chi-square periodogram results of representative animals from each group computed for locomotion (A), feeding (B), RER (C), and energy expenditure (D). Solid black line indicates threshold of significance. For demonstration, the primary peak has been labeled with an open circle, while the secondary peak (if present) has been labeled with an open triangle. **E – H:** Result of chi-square periodogram analysis performed on locomotion (E), feeding (F), RER (G), and energy expenditure (H) across the entire experimental timeline, with the periodicity, in hours, of the two most powerful peaks plotted on the y-axis. Primary (i.e., highest power) peaks are shown as closed circles where color indicates the power of the peak. Secondary peaks are shown as open triangles without color. **Stats:** We performed chi-square periodogram analysis using the zeitgebr::periodogram function in R with a fixed sampling interval and a period search range between 16-32h. The alpha threshold for statistically significant periodicity was set at 0.001.

### ECD disrupts temporal coordination between feeding and RER independent of CB1r signaling

We next asked how ECD altered the relationship of behaviors with respect to one another, rather than to the light/dark environment, in both WT and CB1r KO mice. We performed cross correlation analysis between two continuous physiological measures, allowing us to determine whether changes in one measure were associated with changes in another measure at different time offsets (“time lags”). We assessed the association between feeding and RER (Figures 5A and 5B) to understand how animals tuned their macronutrient oxidation (i.e., RER) in response to nutritional status. As a negative control, we assessed the correlation between energy expenditure and locomotion (Figures 5C and 5D) which, according to the first law of thermodynamics, should remain highly correlated. Cross correlation analysis revealed that the maximum correlation between feeding and RER occurred after 1hr, meaning that a change in feeding was positively correlated with a change in RER 1hr later (Figures 5E and 5F), whereas maximal correlation between locomotion and energy expenditure was immediate, occurring within the same 1hr bin (Figures 5G and 5H). Importantly, ECD significantly reduced the maximum correlation between feeding and RER in both WT and CB1r KO mice, while having no effect on the maximum correlation between locomotion and energy expenditure (Figures 5I-5K). Thus, ECD reduced the temporal coordination between feeding and RER in both WT and CB1r KO animals.

**Figure 5:**
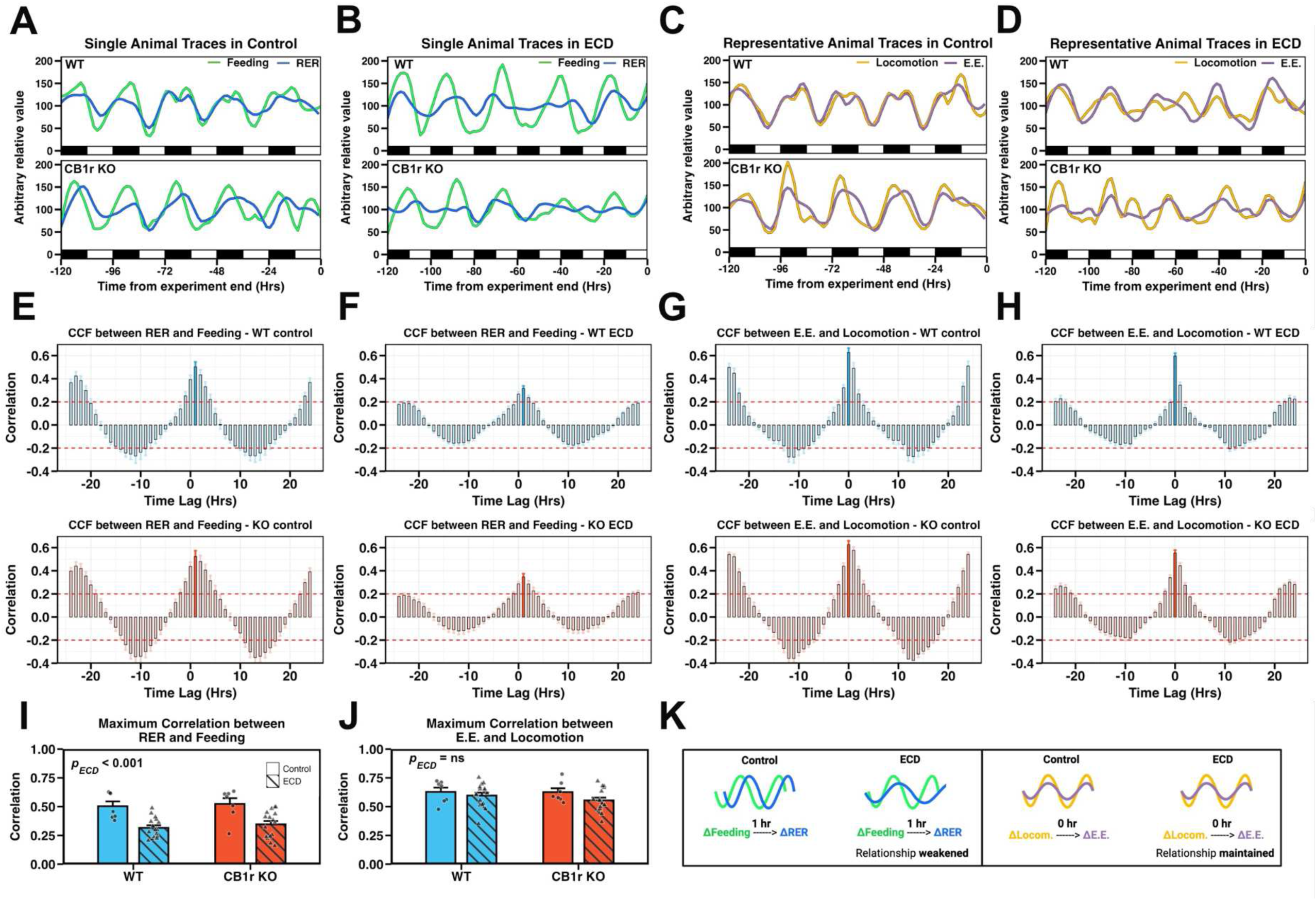
ECD disrupts temporal coordination between feeding and RER independent of CB1r signaling. **A – D:** The final 120hr of experimental feeding and RER (A and B; green and blue lines, respectively) or locomotion and energy expenditure (C and D; orange and purple lines, respectively) of representative WT and KO (upper and lower panels, respectively) male mice in Control and ECD conditions. **E – H:** Analysis of temporal correlation between RER and feeding (E, F) or energy expenditure and locomotion (G, H) in Control and ECD conditions. Time lag, plotted on the x-axis, is the time, in hours, that a change in variable X (RER or energy expenditure) lags a change in variable Y (feeding or locomotion). Maximum correlations are indicated by solid bars. *n*_Control_ = 7/genotype, *n*_ECD_ = 16/genotype. **I and J:** Maximum correlation values between RER and feeding (I; *p*_Int_ = 0.886, ; *p*_ECD_ < 0.001) and energy expenditure and locomotion (J; *p*_Int_ = 0.525, ; *p*_ECD_ = 0.103) for all animals in Control and ECD conditions. Maximum correlation between RER and feeding occurred at time lag = 1, while maximum correlation between E.E. and locomotion occurred at time lag = 0. *n*_Control_ = 7/genotype, *n*_ECD_ = 16/genotype. **K:** Diagram summarizing our finding that ECD weakens the maximum correlation between RER and feeding but does not alter the maximum correlation between energy expenditure and locomotion. **Stats:** Bar plots are expressed as mean +/- standard error of the mean. (I, J) Effect of genotype, experimental condition, and their interaction was assessed using a two-way ANOVA.

### Hepatocyte-specific CB1r is necessary for hormonal disruption, but not weight gain, caused by ECD

The similar behavioral response to ECD between WT and CB1r KO mice, as well as prior literature documenting an importance for hepatic CB1r in metabolic regulation^16,17^, led us to ask if hepatic CB1r signaling was necessary for ECD-induced metabolic disruption. To test this, we exposed male albumin-Cre^+/+^ CB1r^fl/fl^ (L-CB1r KO) mice, which lack CB1r in hepatocytes, and WT littermates to Control and ECD conditions. We found that while L-CB1r KO was insufficient to protect against the weight gain effect of ECD (Figure 6A), these mice were protected against increases in plasma leptin and insulin (Figures 6B and 6C), with a nearly significant interaction for rescued glucagon levels (Figure 6D). L-CB1r also mitigated the increase in plasma triglycerides caused by ECD (Figure 6E), but there was no effect of ECD or genotype on hepatic triglyceride levels (Figure S3D). ECD elevated levels of PYY and amylin in both genotypes (Figure S3B). Finally, ECD had no effect on inflammatory proteins MCP1 and TNF- α, but there was a trend toward elevated IL-6 in WTs, but not L-CB1r KOs (Figure S3B).

**Figure 6:**
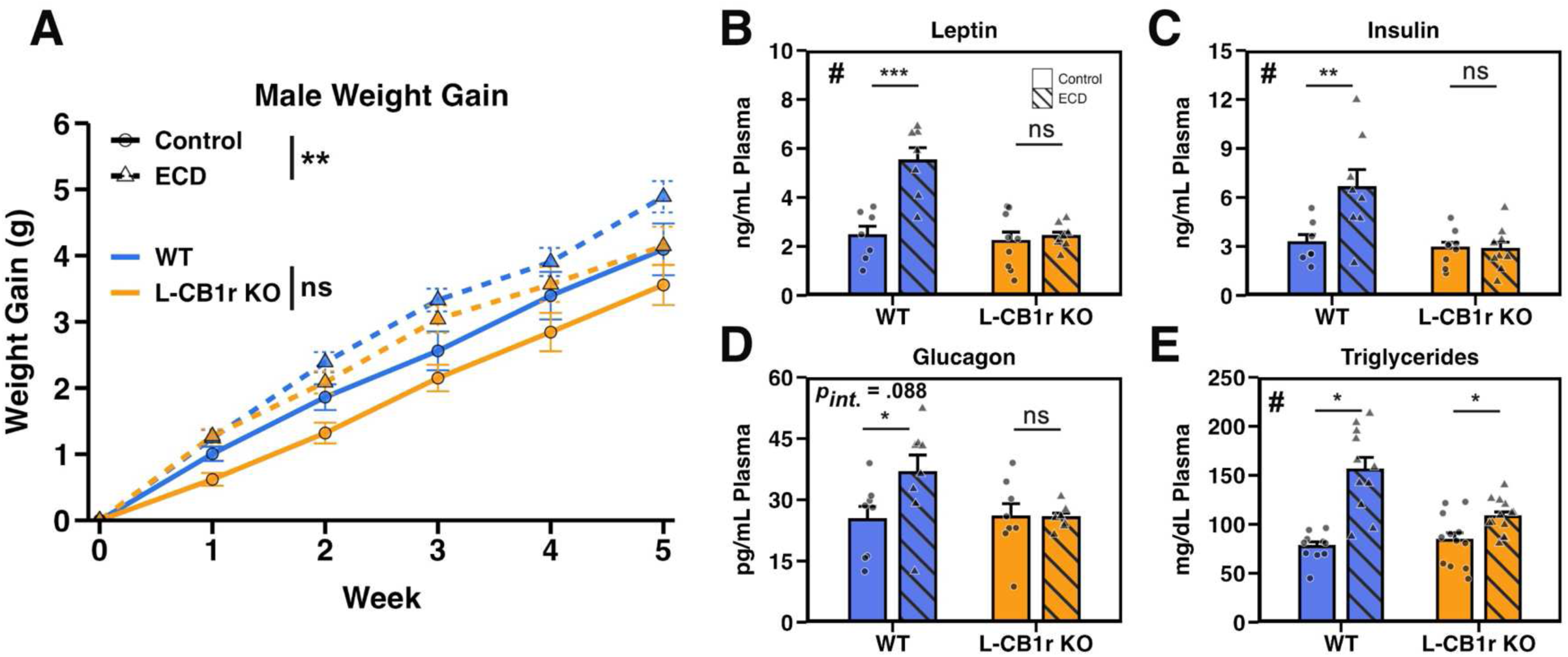
Hepatocyte-specific CB1r is necessary for hormonal disruption, but not weight gain, caused by ECD. **A:** Weight gain of male mice recorded manually every 7 days in Control and ECD conditions. Data were analyzed using a linear mixed effects model (*p*_Genotype x Condition x Week_ = 0.451) and within-group follow-up analysis of the effect of ECD performed using estimate of marginal means (*p*_WT_ = 0.049; *p*_KO_ = 0.017). *n*_Control_ = 16-18/genotype/experimental condition/week. *n*_ECD_ = 26-28/genotype/experimental condition/week. **B – E:** Plasma levels of leptin (B; *p*_Int_ < 0.001, *p*_WT_ = 0.700 *P*_KO_ < 0.001), insulin (C; *p*_Int_ = 0.013, *p*_WT_ = 0.926 *P*_KO_ = 0.002), glucagon (D; *p*_Int_ = 0.088, *p*_WT_ = 0.970 *P*_KO_ = 0.018), and triglycerides (E; *p*_Int_ = 0.003, *p*_WT_ = 0.030 *P*_KO_ = 0.039) from male mice collected at ZT 12. Significant interaction between genotype and ECD denoted by “#”, significant post-hoc t-test indicated by “*”. *n* = 7-13/genotype/experimental condition. **Stats:** Data are reported as mean +/- the standard error of the mean. For weight gain (A), main effects and follow up comparisons were performed using a mixed effects model and estimate of marginal means, respectively (see methods). For plasma metabolite assessment (B-E), we performed a two-way ANOVA (or non-parametric ART ANOVA if assumptions were not met; see methods). Following a significant interaction term, follow-up post-hoc t-tests were performed and *p*-values adjusted using the Bonferroni method. \**p* < 0.05, \*\**p* < 0.01, \*\*\**p* < 0.001.

### ECD remodels hepatic metabolic transcriptional profiles in a CB1r-dependent manner

We sought to define how ECD affects liver function on the transcriptional level, and any role that CB1r may play in mediating this. To accomplish this, we employed NanoString Metabolic Panel gene expression analysis on whole liver tissue from WT and L-CB1r KO males. We first assessed functional gene sets that were oppositely regulated by ECD in WT vs L-CB1r KO mice (Figure 7A; complete gene set results in Figure S6A). In WT males, ECD upregulated gene pathways involved in mitochondrial respiration, reactive oxygen response, and DNA and cellular damage (Figure 7A). Remarkably, these pathways were downregulated by ECD in L-CB1r KO males.

**Figure 7:**
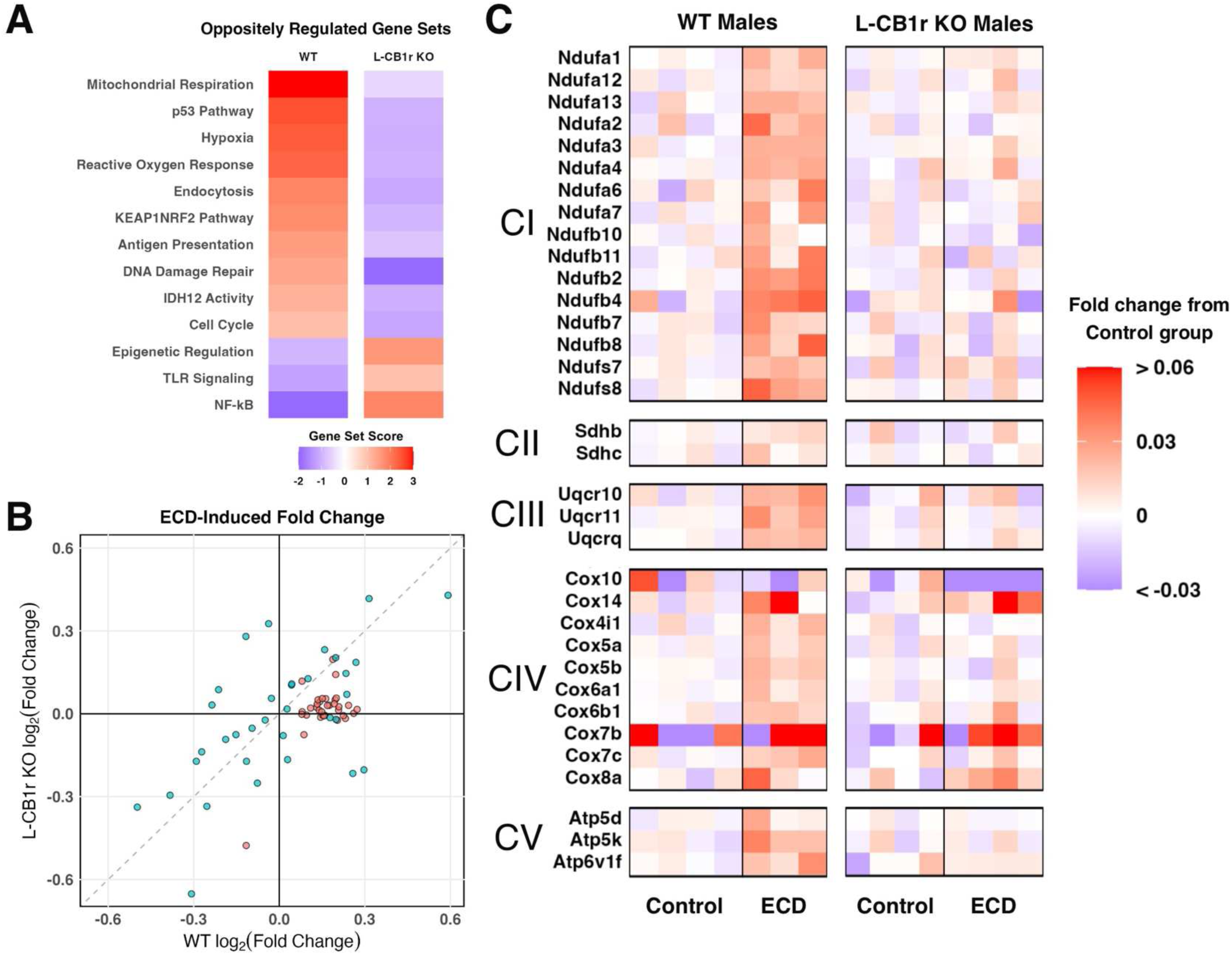
ECD remodels hepatic metabolic transcriptional profile in a CB1r-dependent manner. **A – C:** Results of NanoString nCounter Metabolic Pathway, showing the effect of ECD on gene expression in WT and L-CB1r KO mice separately. A) Gene sets for which the difference in gene set score between WT and L-CB1r KO was greater than |2|. Presented in descending order of gene set score for WT mice. B) Scatterplot showing log_2_(fold change) of individual transcripts belonging to the Mitochondrial Respiration gene set. C) Heatmaps of relative expression of gene transcripts involved in the electron transport chain. Transcripts are separated into and labeled by the super-complex they comprise or facilitate (CI – CV). Within each genotype, individual samples are expressed as fold change with respect to the mean of the within-genotype Control group.

Mitochondrial respiration was the most upregulated gene set in WT animals, and disruption to this process could likely account for the upregulation of other gene sets, such as reactive oxygen response, cellular stress, and DNA damage. We assessed individual gene expression to understand which components of mitochondrial respiration were upregulated by ECD (Figure 7B). We observed that a cluster of genes that were consistently upregulated in WT mice following ECD encoded components of the electron transport chain, the final step in mitochondrial respiration. Additional pathway analysis confirmed that ECD upregulated nearly all genes in this pathway in WT animals, while there was no effect in L- CB1r KOs (Figure 7C). Thus, ECD altered the cellular metabolic transcriptional profile of the liver in a CB1r-dependent manner.

## Discussion

Circadian misalignment is driven by behavioral and environmental factors, such as shift work or jet lag, and is common in modern society^5^. Yet, the pathways by which circadian misalignment leads to metabolic dysregulation remain incompletely defined. Using a model of ECD, we aimed to investigate if the endocannabinoid system, which is known to be important in metabolic regulation, may represent a possible pathway that links circadian misalignment and metabolic changes. Our results suggest that CB1r may be dysregulated under ECD via an emergent coordination of AEA and 2-AG levels both systemically and in the liver. We also showed that ECD induced weight gain, dyslipidemia, and metabolic hormone disruption in a CB1r-dependent manner. Curiously, this metabolic protection could not be attributed to a different behavioral response to ECD in global CB1r KO, suggesting a potential peripheral mechanism. We showed that L-CB1r KO was sufficient to protect against much of the metabolic phenotype induced by ECD. This observation may be explained by the fact that ECD altered liver mitochondrial respiration in WT, but not L-CB1r KO, males.

We have previously shown that 10-week exposure to ECD leads to significant metabolic consequences in male mice^19^. In this study we used a shorter 5-week ECD challenge to assess the relationship between early behavioral and physiological changes and metabolic disruption. In males, ECD increased weight gain in WT but not CB1r KO mice; in females, ECD did not increase weight gain in either genotype. This is unsurprising, since female mice are more resilient than males in the face of certain metabolic challenges, such as high-fat diet^22^. Detailed future studies will be needed to understand the nature of this sex difference.

Both total levels and daily rhythms of endocannabinoids are altered in individuals with obesity^12–14^, but whether such changes are a cause or consequence of metabolic disruption is unclear. We assessed endocannabinoid levels at this early stage of metabolic disruption to minimize the influence of increased body weight, which is exacerbated over longer exposures to ECD^19^ and is known to influence the level of circulating endocannabinoids^12^. While we did not observe an overall change in AEA or 2-AG levels in plasma or liver following ECD, we did observe that these ligands became significantly and positively correlated in ECD. Similar observations in female WTs, which did not exhibit excessive weight gain, indicate that this phenomenon is not simply a result of increased body weight, but may instead be a contributing factor to weight gain. The emergent coordination of endocannabinoids is relevant because the CB1r activation profile differs in the presence of both AEA and 2-AG compared to either ligand alone^23,24^. Therefore, a novel coordination in endocannabinoid levels could potentially dysregulate CB1 activity. This observation invites future studies to test whether coordinated endocannabinoid levels, without a change in overall concentrations, are necessary and/or sufficient to elicit metabolic disruption.

The ECD-induced metabolic phenotype in male mice occurred without a detectable change in gross energy intake or expenditure. This led us to carefully analyze the temporal coordination of these behaviors and their impact on physiology. Using continuous long-term behavioral phenotyping, we determined that ECD led to weakened and imprecise physiological rhythms that were often misaligned with environmental cues. These observations reinforce that the timing, not just the quantity, of energy intake and expenditure can influence metabolic homeostasis. We additionally showed that locomotion, RER, feeding, and energy expenditure were similarly disrupted by ECD, implicating the disruption of these processes in contributing to the metabolic phenotype. However, and somewhat surprisingly, our phenotyping data did not provide evidence of a behavioral basis for the observed differences between WT and CB1r KO mice. In other words, even though CB1r KO mice were protected from the development of the weight-gain and metabolic hormone phenotype, their behavioral responses to ECD were indistinguishable from their WT counterparts.

We observed that ECD weakened the temporal correlation between feeding and RER, which suggests inefficient metabolic response to nutritional cues. Again, this could be both a cause and consequence of weight gain and metabolic hormone disruption. For example, if ECD results in frequent misalignment between nutrient intake and endogenous metabolic hormone rhythms, metabolic response to nutritional status could worsen and lead to metabolic disruption. Alternatively, pre-existing metabolic conditions such as insulin resistance would plausibly diminish the ability for feeding to elicit carbohydrate oxidation and subsequent RER elevation. Yet, both WT and CB1r KO males exhibit a weakened relationship between feeding and RER following ECD, despite CB1r KOs being protected from the metabolic phenotype. This supports the notion that inefficient nutrient handling precedes ECD-induced weight gain and hormone disruption, and that CB1r signaling is necessary to translate such metabolic inefficiency into this phenotype.

Altered hepatic function is frequently observed in different models of systemic metabolic dysfunction^25^. The liver is also highly regulated on a circadian timescale, showing significant daily changes in clock gene expression both *in vivo*^18^ and *ex vivo*^26^, as well as a high degree of daily regulation of the transcriptome^27^ and proteome^28^. Our study contributes to a growing body of research implicating hepatic endocannabinoid signaling in regulating metabolic function by showing that hepatocyte CB1r is necessary for hormonal disruption and dyslipidemia, but not weight gain, following ECD.

To understand how ECD affects the transcriptional profile of the liver, and any mediating role of CB1r in this process, we performed gene transcription analysis on WT and L-CB1r KO livers. We theorized that coordinated endocannabinoid levels following ECD may lead to a differential pattern of CB1r activity. Given that CB1r activation is known to inhibit mitochondrial oxidative phosphorylation through multiple pathways^29–31^, this could lead to cellular dysfunction including ROS accumulation and pseudo-hypoxic conditions. In support of this mechanism, transcripts related to mitochondrial respiration were highly upregulated in WT mice following ECD, possibly to compensate for ECD-induced inhibition of oxidative phosphorylation. Furthermore, ECD upregulated transcripts related to reactive oxygen response, DNA damage, and hypoxia response gene sets, all of which could potentially result from dysfunctional mitochondrial respiration^32,33^. This is especially interesting in light of prior research showing that hepatic mitochondrial dysfunction precedes obesity, insulin resistance, and dyslipidemia^34^.

While CB1r activation can suppress mitochondrial respiration through multiple mechanisms, its action on the FoxO1 pathway stands out due to its connection with insulin signaling. CB1r activates the insulin-sensitive FoxO1 signaling pathway, which inhibits mitochondrial respiration^35,36^. If hepatic CB1r is overactivated following ECD (possibly via coordinated endocannabinoid levels), we may expect a compensatory increase in insulin to inhibit FoxO1 signaling, as well as an upregulation of electron transport chain transcripts to counteract the FoxO1-mediated suppression of oxidative phosphorylation. This could explain why ECD elevates plasma insulin in WT, but not L-CB1r KO males, and should be interrogated in future studies.

Understanding the links between metabolic dysregulation and the environmental light/dark cycle is critical. Genetic models of circadian clock disruption have clearly demonstrated links between clock function and metabolism in a variety of metabolically important tissues, such as liver^37^, pancreas^38^, adipose tissue^39^, and skeletal muscle^40^. These models have been important to build an understanding of the molecular mechanisms linking circadian processes and metabolic function. However, genetic circadian disruption is not a common occurrence in humans, and as such, environmental models of disrupted circadian timing would seem to have greater translational relevance. Many models of environmental circadian desynchronization, from repeated phase shifts^41,42^, to light at night^43,44^, to altered environmental cycle lengths^45^ (as in the present study) have been used to demonstrate the impact of dysregulated timing on metabolism. These models each have their own strengths and weaknesses, but they do provide ample evidence that disrupting the light/dark cycle in which an organism lives has severe biological consequences. Our study provides evidence that the endocannabinoid system acts as a link between circadian biology and metabolic physiology.

The role of the endocannabinoid system in obesity and metabolic function is a very active area of research^6^, and excellent work in humans has linked obesity to changes in the diurnal rhythms of circulating endocannabinoids^14^. Our finding that disruption of the endocannabinoid system globally can block the metabolic impact of circadian desynchronization without an impact on behavior has important implications for development of potential interventions. Further, demonstrating that many key aspects of metabolic disruption are related to hepatic endocannabinoid signaling may lead to development of more personalized approaches to treating the effects of chronic circadian disruption.

Disruptions to human circadian rhythms, such as those caused by shift work, light at night, and social jet lag are pervasive in society and are likely to be a permanent fixture in the future. As industrialization continues to separate us from the photic environment we evolved to live in, the negative impact on our biological clocks will lead to even greater chronic health risks. While increasing acknowledgment of these negative effects may result in better work schedules and exposure guidelines, understanding cellular and molecular links between circadian disruption, behavioral and physiological misalignment, and metabolic dysregulation will provide us with potential countermeasures to some of these impacts when behavioral mitigation is not possible.

## Methods

### Mouse lines

The CB1r^−/−^ global and liver-specific (L-CB1r KO; Alb-Cre^+/+^ CB1r^f/f^) mice used in this study were originally generated on a C57/Bl6J background^16,17^ and subsequently backcrossed to a C57/Bl6N background for at least 10 generations. Global CB1r WT and KO mice were generated by breeding heterozygous mice. L-CB1r^−/−^ mice were generated by crossing mice homozygous for the CB1r-floxed allele (CB1r^f/f^; Jackson Laboratory stock number 36107^46^), which were on a predominantly C57BL/6N background (seven to eight crossings), with mice expressing the bacterial Cre recombinase driven by the mouse albumin promoter (triglyceride[Alb-cre]’21 Mgn) that had been backcrossed seven times to a C57BL/6J background (Jackson Laboratory stock number 3574^47^) to obtain CB1r^f/f^ × CB1r^f/f^ Alb-Cre breeding pairs (thus a mixed C57BL/6J and 6N background).

Mice were genotyped using DNA extracted from ear biopsies as described^48^. The CB1r KO allele is identified by PCR using the Amfisure PCR Master mix (2X) from GenDepot (cat# P0311-025) and the following primers: CB1 common CTC CTG GCA CCT CTT TCT CAG TCA CG; CB1 knockout TCT CTC GTG GGA TCA TTG TTT TTC TCT TGA and CB1 wildtype TGT GTC TCC TGC TGG AAC CAA CGG with bands at 284 bp for WT and 334 bp for KO. The Albumin Cre transgene is identified by PCR using the following primers: Common TTG GCC CCT TAC CAT AAC TG, wild type forward TGC AAA CAT CAC ATG CAC AC and mutant forward GAA GCA GAA GCT TAG GAA GAT GG with bands at 351 bp for transgene negative and 390 bp for transgene positive.

### Animals and Housing Conditions

All animal procedures and experiments were approved by the University of Massachusetts – Amherst Institutional Care and Use Committee. Global CB1r WT and KO littermate mice, aged 7-8wk old, were randomly assigned to either be group housed in standard mouse cages (*n* = 2-5/cage, same sex and genotype within cage) or to be singly housed in the TSE Phenomaster system (TSE-Systems International, Berlin, Germany) for behavioral phenotyping (and described more fully below). Animals were allowed 1wk to acclimate to their new environment before experimental conditions and recording began at 8-9wk of age. Control or ECD conditions lasted for 5 weeks. Mice housed in the TSE Phenomaster system were collected at Zeitgeber Time (ZT; defined further, below) 12 (lights off), while group-housed mice were collected at one of the following time points: ZT 0, 4, 8, 12, 16, 20. Hormone and lipid assays were performed using plasma or liver collected from global WT and CB1r KO mice housed in the TSE and collected at ZT 12. Endocannabinoid levels were measured using plasma collected from group-housed animals collected at various time points.

L-CB1r WT and KO littermate mice, aged 8-9wk old, were randomly assigned to be singly housed either in standard mouse cages or in the TSE Phenomaster system. They were allowed 1wk to acclimate before experimental conditions and recording began at 9-10wk of age. Control or ECD conditions lasted for 5 weeks. All mice were collected at ZT 12 and utilized for plasma and liver hormone analysis. Liver tissue was used for NanoString gene expression analysis.

All mouse cages were placed in ventilated, light-tight boxes (Actimetrics Circadian Cabinets; Actimetrics, Wilmette, IL, USA) that could be programmed to control light settings (ClockLab Chamber Control; Actimetrics, Wilmette, IL, USA). All animals had ad-lib access to food (Prolab Iso-Pro RMH 3000, LabDiet Cat# 5P76) and water. Cage changes were performed every 7 days, at which time body weights were also recorded. Light intensity was measured using a lux meter (Dr. meter, cat# LX1010B) and determined to range between 40- 50 lux, as measured from within a fully assembled, closed cage. Temperature within the light-tight boxes was maintained between 22-25°C.

### Tissue Collection

Mice were rapidly decapitated without anesthesia, and trunk blood was immediately collected and placed into a blood collection tube containing EDTA (Monoject EDTA (K3) 15%; Covidien, Dublin, Ireland) which was placed on ice while tissue collection continued. A liver sample (right medial lobe) was collected and placed in 1.5mL Eppendorf tubes, placed in powdered dry ice to freeze, and subsequently stored long term at -80°C. Following tissue collection, tubes containing trunk blood were centrifuged at 1,300 x g for 15min at 4°C to separate plasma from serum. The plasma (top layer) was then isolated via pipette and placed into 1.5mL Eppendorf tubes on powdered dry ice to freeze and stored long-term at - 80°C until analysis.

### Defining *Zeitgeber* Time (ZT)

*Zeitgeber* time is a method of expressing time relative to the lighting cues of an environment. By convention, one full lighting period is expressed as 24 “ZT Hours”, where ZT 0 is defined as the beginning of lights on. Under Control conditions, (12hr light / 12hr dark) lights come on at ZT 0 and, 12 hours later, lights go off at ZT 12. In the ECD model used in this study (10hr light / 10hr dark) lights come on at ZT 0 and, 10 hours later, lights go off at ZT 12. Under this model, each ZT hour contracts to become 50-minutes long, so that the entire 20hr period can be expressed as 24 “ZT hours”.

### Behavioral and physiological analysis

Locomotion, feeding, drinking and indirect calorimetry were measured using the TSE Phenomaster system. Locomotion was tracked using infrared beam breaks. In locomotor analyses, we included total locomotion in the X and Y axes of the cage, but not vertical locomotion (e.g., rearing). Food and water consumption were measured with precision suspension-scales controlled by the Phenomaster software. Food baskets were equipped with a collection dish to minimize the effect of food spillage. To calculate energy intake, the metabolizable energy content of the standard chow diet (Prolab Iso-Pro RMH 3000, LabDiet Cat# 5P76) was used, which equates to 3.16 Kcal/g food. RER and energy expenditure were calculated via indirect gas calorimetry, in which air was sampled from each cage for ∼3.5 minutes, once per hour, to attain reading of VO2 and VCO2. Gas detection device was calibrated weekly using a set of three, highly precise spec gasses.

### Behavior peak detection algorithm

To compute behavioral peaks (Figure 3), we developed an algorithm in Python that ran a convolution with a Gaussian kernel (similar to a moving average, which would be a convolution with a uniform kernel). First, we binned the data into 5-minute bins, then ran a convolution to smooth the data. We then identified activity minima as the point of lowest activity within a certain sampling period for each point. Then we defined peaks as the maximal point of activity between each two minima. We discarded any peaks within 12 hours of each other. For each window between two peaks, the onset was defined as the last datapoint below the β^th^ percentile of points in that window. The model parameters for beta, the sizes of the sampling windows, and the gaussian kernel were determined empirically by modeling the algorithm to the behavioral data from mice under Control conditions, then applying these same parameters to mice under both ECD and Control conditions.

### Periodogram Analysis

Periodograms were generated using the “Rethomics” package^49^ with a fixed sampling interval and a period search range between 18-30h. Analysis was performed over the entire 35-day experimental condition. Statistical significance of rhythmic periods was evaluated using a chi-squared test with a alpha threshold of 0.001.

### Cross Correlation Analysis

Cross correlation analysis (Figure 5) was performed on RER and feeding or energy expenditure and locomotion, which were binned into 1hr bins but otherwise not transformed. Analysis was performed using Stats::ccf function in R. To visualize these physiological measures on the same scale (Figures 5A-5D), and to account for between- animal differences in these measures, representative data were first transformed as percent change from the 120hr mean, within measure. Then, to account for differences in the magnitude of daily fluctuations between measures, each measure was multiplied by an empirically determined measure-specific constant so that the amplitudes of feeding and RER or locomotion and E.E. were comparable.

### Data exclusion criteria

Mice periodically exhibited food hoarding behavior by removing a large amount of food and allowing it to waste on their cage floor, complicating certain feeding analyses. To minimize the effect of this on quantitative feeding analysis, we employed outlier replacement as follows: feeding was first binned into daily light and dark phases, and means were computed across the entire experiment within group (genotype x experimental condition x phase). Then, any binned feeding value which exceeded Q3 + 2.5 * IQR of its group was replaced with the median value from that group, where Q3 is the third quartile and IQR is the interquartile range. This significantly reduced righthand skewedness in feeding data caused by this hoarding behavior and resulted in the replacement of 4.6% of data points from the dataset. RER values less than 0.7 or greater than 1.2 were excluded from analysis, which accounted for <0.001% of data points. Such values are likely the result of technical, rather than biological, anomalies. Analysis-specific outlier exclusion, if performed, is described in the respective sections below.

### Triglyceride Assay

Triglyceride content in the plasma and liver was assessed using the Triglyceride Colorimetric Assay Kit (Cat no. 10010303; Cayman Chemical Company, Michigan, USA) according to manufacturer’s instructions. For plasma triglyceride quantification, 10uL of plasma was diluted 1:2 with 1X Standard Diluent Assay Reagent. For liver tissue, approximately 25mg of fresh frozen liver tissue was isolated and 25uL of NP40 solution per mg liver tissue was added to the Eppendorf tube containing a 5mm stainless steel bead. Liver tissue was then homogenized for 2min at 35Hz (TissueLyser LT; Qiagen, Hilden, Germany) and subsequently centrifuged at 10,000 x g for 10min at 4°C. Supernatant was then isolated and stored at -80°C until day of assay, at which point 10µL of undiluted sample was added to each well. For plasma triglyceride assessment, 10µL of plasma (diluted 1:2 with Standard Diluent Assay Reagent) was added to each well. Absorbance was read at 540nm using a microplate reader (Biotek Epoch, Winooski, VT, USA) and Gen5 software for data processing. Samples were run across multiple plates, corrected for by using the same calibrator sample ran on each plate. In cases where the %CV between duplicates was greater than 10, the replicate that was more distal from the group (genotype x experimental condition) mean was eliminated. Statistical outliers were identified and removed as follows: duplicate wells for each sample were averaged and means for each metabolite were then computed within group. Then, any value less than Q1 – 1.5 * IQR or greater than Q3 + 1.5 * IQR was excluded from analysis, resulting in an exclusion of 2.4% of datapoints.

### Magpix xMap Metabolic Hormone Assay

Plasma concentration of Amylin, IL-6, Glucagon, TNF-α, Insulin, Leptin, MCP1, GIP, and PYY was quantified using the customizable Milliplex Mouse Metabolic Hormone Expanded Panel and Luminex XMAP technology Magpix (Cat no. MMHE-44K; EMD Millipore, Darmstadt, Germany) according to manufacturer’s instructions. For this assay, we used undiluted plasma derived from trunk blood (see Methods: Tissue Collection) that was treated only with EDTA (no enzyme inhibitors). Samples were run across multiple plates, corrected for by using the same calibrator sample run on each plate. In cases where the coefficient of variance between duplicates was greater than 10%, the replicate that was more distal from the group (genotype x experimental condition) mean was eliminated. Statistical outliers were identified and removed as follows: duplicate wells for each sample were averaged and means for each metabolite were then computed within group. Then, any value less than Q1 – 1.5 * IQR or greater than Q3 + 1.5 * IQR was excluded from analysis, resulting in an exclusion of 5.3% of datapoints.

### Endocannabinoid Quantification

Detection of endocannabinoid concentrations was performed using liquid chromatography / tandem mass spectrometry (LC-MS/MS) as previously described^50,51^. Briefly, frozen tissue samples were quickly weighed and manually homogenized using a glass rod in a borosilicate glass tube containing 2 mL of acetonitrile and 100µl of Internal Standard (5 pmol d8-AEA [Cayman Chemical Company, #390050] and 5 nmol d8-2-AG [Cayman Chemical Company, #362160]). For blood samples, plasma was transferred to glass tubes prepared in the same manner directly into the acetonitrile. Prepared samples were sonicated for 20 minutes before incubating overnight at -20°C to precipitate proteins. The next day samples were centrifuged (1,800 rpm at 4°C for 3–4 min) to remove particulates, the supernatant transferred to a new glass tube, and tubes evaporated under nitrogen gas. The tube sidewalls were washed once with 250μl acetonitrile to recollect any adhering lipids and completely evaporated again under nitrogen gas. Finally, samples were resuspended in 200μl of acetonitrile before storage at -80°C until analysis by LC-MS/MS. Analyte (AEA and 2-AG) concentration (in pmol/mL for plasma or pmol/g for liver tissue) were normalized to sample weight. Statistical outliers were identified and removed as follows: means were computed within group (sex x ZT x experimental condition x endocannabinoid). Then, any value less than Q1 – 1.5 * IQR or greater than Q3 + 1.5 * IQR was excluded from analysis, resulting in the removal of 6.6% of datapoints.

### RNA Extraction for NanoString Analysis

Following removal from -80°C, 5mg of mouse liver were collected into a tube containing RLT buffer (Qiagen). Liver tissue was disrupted in the TissueLyser LT by rapid agitation (2 minutes at 50 Hz) in the presence of 5mm steel beads. RNA was extracted using the RNeasy micro kit (Qiagen cat#74004) following the manufacturer’s instructions. Extracted RNA concentration/quality was analyzed using Qubit and a Nanodrop Spectrophotometer prior to NanoString analysis.

### NanoString Gene Expression Analysis

Transcriptional analysis of liver RNA samples was performed using the nCounter Metabolic Pathways Panel by NanoString. The nCounter panels were run at the Center for Personalized Cancer Therapy (CPCT) Genomics Core at UMass Boston according to supplier protocols. Analysis of NanoString data was performed as previously described^52^, with exceptions stated below.

Quality control measures of all samples fell within the following ranges: Imaging QC =0.98 – 1.0; Binding Density = 1.03 – 1.47; Positive Control Linearity = 0.99 – 1.0; Field of View = 96 – 100%; Limit of Detection = 13.17 – 24.63. Because samples were run over two different plates with identical batch IDs, inter-plate calibration was not necessary. As recommended by the manufacturer, neither background subtraction nor background thresholding were performed. Between-sample normalization was performed via a two-step process in the Advanced Analysis (Version 2.0.134) module within nSolver using default settings, as described in the nSolver 4.0 Analysis Software User Manual. This automatically chose optimal normalization probes for each comparison being performed by removing the most unstable housekeeping genes. For comparison of Control vs ECD in WT animals, the selected housekeeping genes for normalization were: *Sdha, Cog7, Oaz1, Mrps5, Fcf1, Sap130, Dhx16, Dnajc14, Abcf1, Ubb,* and *Usp39.* For comparison of Control vs ECD in L- CB1r KO animals, this list included all the genes listed for WTs, plus *Tlk2* and *Edc3*.

Gene set analysis (GSA) was also performed in nSolver using the Advanced Analysis module. GSA is an averaging of the differential expression measures of all genes within a given pathway (gene set). We computed “Directed” Global Significance Score, which accounts for both magnitude and direction of differential expression, as described by NanoString. A high gene set score indicates that a majority of genes in the gene set exhibit changes in expression, in the same direction, across multiple samples.

### Statistics

All data processing and analysis were performed in R^53^ using the RStudio environment^54^, except for the in-house peak and onset detection algorithm which was created in Python.

For 2-way ANOVAs (Figures 1D-G, 3I-L, 5I & J, 6B-E, S1A-H, and S3A-D), data were first tested against the assumptions of normality of residuals and homoscedasticity using the Shapiro-Wilk test and Levene’s test, respectively. If assumptions were met, group differences were assessed using a standard ANOVA with type III sums of squares. If either assumption was violated, group differences were assessed using the aligned rank transform (ART) procedure to allow for nonparametric analysis of factorial designs while preserving interaction effects. When a significant three-way interaction term was present, the effect of ECD was tested within each genotype by performing estimated marginal means, applying Bonferroni correction for multiple testing.

Phase-specific behavioral and physiological measures (Figures 2C-F) were first tested against the assumptions of normality of residuals and homoscedasticity using the Shapiro-Wilk test and Levene’s test, respectively. If assumptions were met, group differences were assessed using a linear mixed effects model including genotype, experimental condition, and phase as fixed effects with animal ID as a random intercept to allow for repeated measures across phase. If either assumption was violated, group differences were assessed using ART ANOVA on the same model structure. Planned t-tests were performed comparing the average activity of in the light versus dark phase, within genotype and experimental condition, applying Bonferroni correction for multiple testing.

For longitudinal, repeated measures such as weight gain or energy balance (Figures 1C, 2AC B, 6A, S2A, and S4A) the assumptions of normality of residuals and homoscedasticity were verified visually due to large sample size, which is known to yield overly sensitive Shapiro-Wilk and Levene’s Test results. All datasets met assumptions and were analyzed using linear mixed effects model (lmerTest::lmer in R) with genotype, experimental condition, and time (continuous) as fixed effects with animal ID as a random intercept to allow for repeated measures. Fixed-effect significance was tested using Type III ANOVA with Satterthwaite’s method for degrees of freedom. Planned estimate of marginal means to test the effect of ECD on weight gain over time within genotype was performed (emmeans::emmeans function in R), applying Bonferroni correction for multiple testing.

For endocannabinoid correlation analysis (Figures 1A CB and S1I & J) the relationship between AEA and 2-AG was evaluated using Pearson’s correlation. Pearson’s correlation coefficient (r) was reported as a measure of effect size, along with the corresponding *p*-value to assess statistical significance.

## Supporting information

Supplemental Tables - Statistics Results

## Resource availability

### Lead contact

Further information and requests for resources and reagents should be directed to and will be fulfilled by the lead contact, Ilia Karatsoreos.

### Materials availability

This study did not generate new unique reagents.

### Data and code availability

All data reported in this paper will be shared by the lead contact upon request.

All original code will be made available by the time of publication of this manuscript.

Any additional information required to reanalyze the data reported in this paper is available from the lead contact upon request.

## Declaration of generative AI and AI-assisted technologies in the writing process

During the preparation of this work the authors used ChatGPT in order to improve readability and succinctness of parts of the Methods section. AI was not used in the writing of the summary, introduction, results, or discussion. After using this tool/service, the authors reviewed and edited the content as needed and take full responsibility for the content of the published article.

## Author Contributions

Conceptualization, I.K., N.B., D.P.; Funding Acquisition: I.K.; Methodology, B.B., N.B., D.P.; Project Administration: B.B.; Investigation, B.B., G.D., S.A., J.W., G.P., C.H., W.S.; Data Curation, B.B., G.D.; Formal Analysis, B.B.; Visualization, B.B.; Supervision: I.K., M.H.; Writing – First Draft, B.B., I.K.; Writing – review and editing, B.B., G.D., S.A., J.W., G.P., C.H., M.H.

## Acknowledgements

The authors would like to thank Tanya Leise, PhD, who lent her uniquely expert opinions on analysis of periodic data. We would also like to thank Mary Harrington, PhD and Cecelia Hillard, PhD, for their input on early versions of this manuscript. We thank Krista Gile, PhD, and Anna Liu, PhD, at UMass Amherst Department of Mathematics and Statistics for their input on statistical analysis.

