## Supplementary figures and images for "Endocannabinoid signaling is a critical link between circadian desynchronization and metabolic dysfunction"

### Supplemental Figures

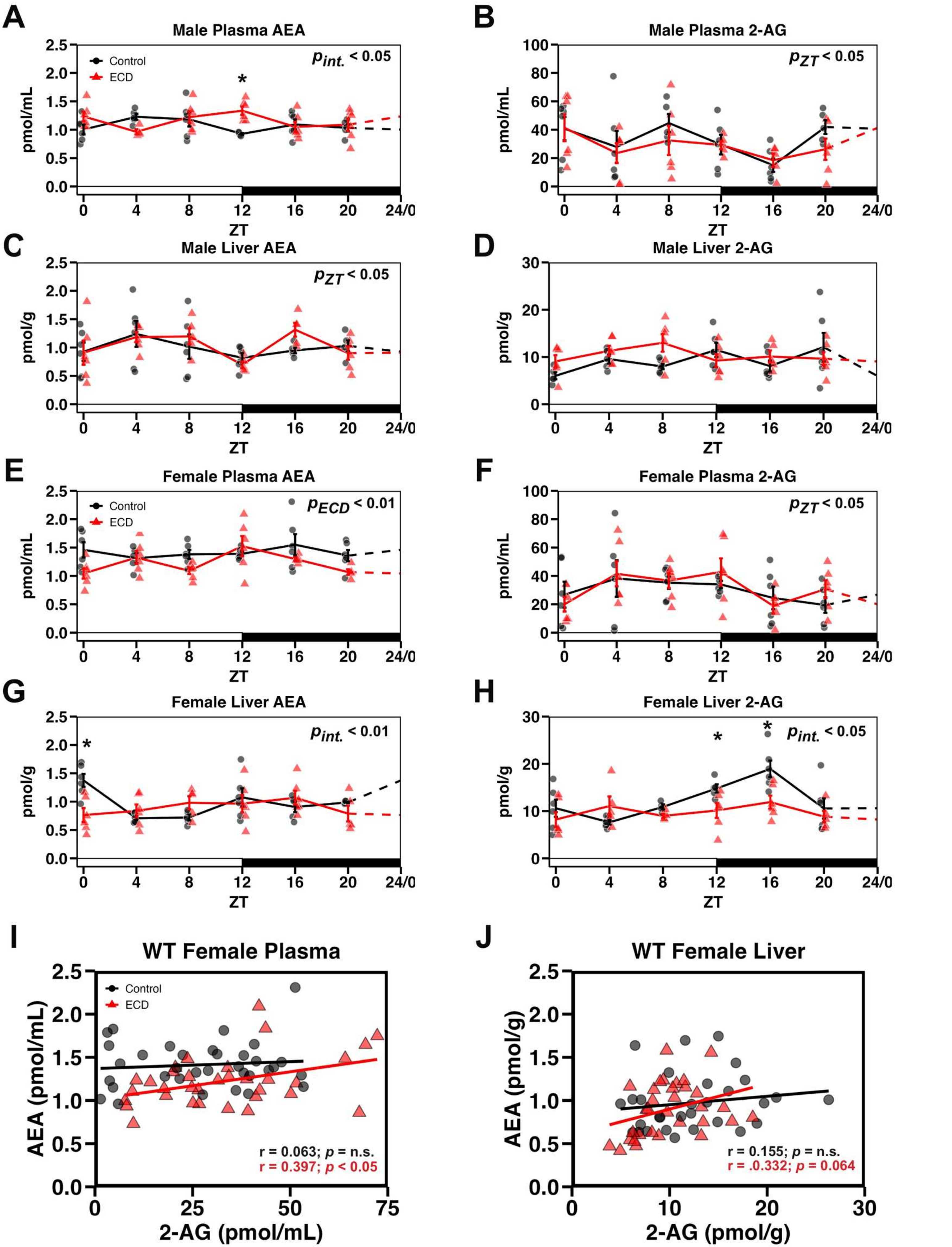

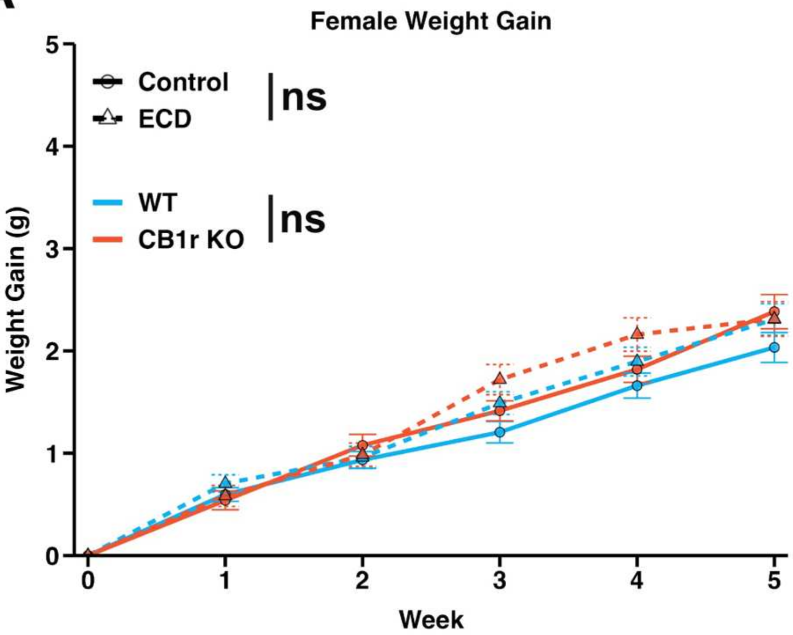

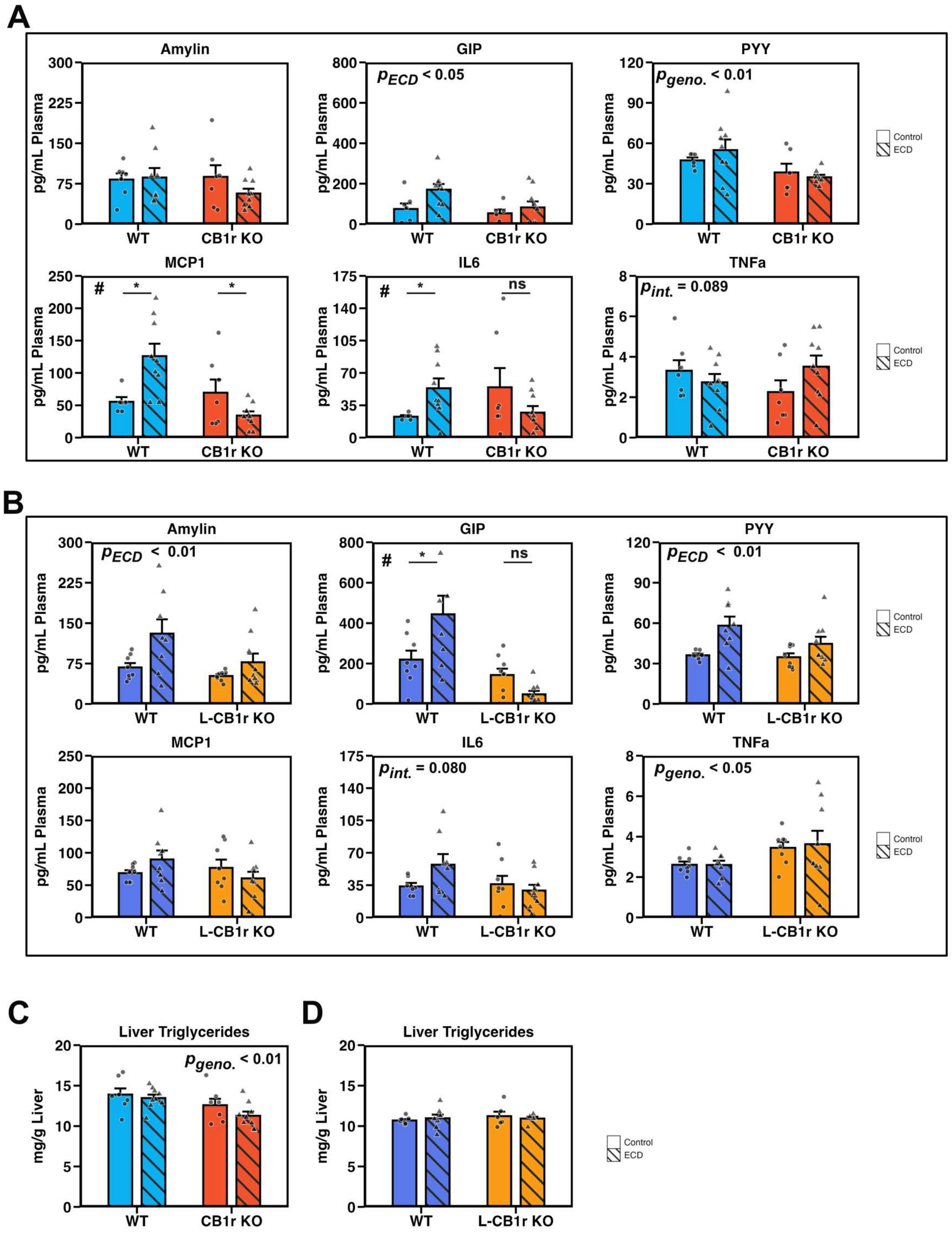

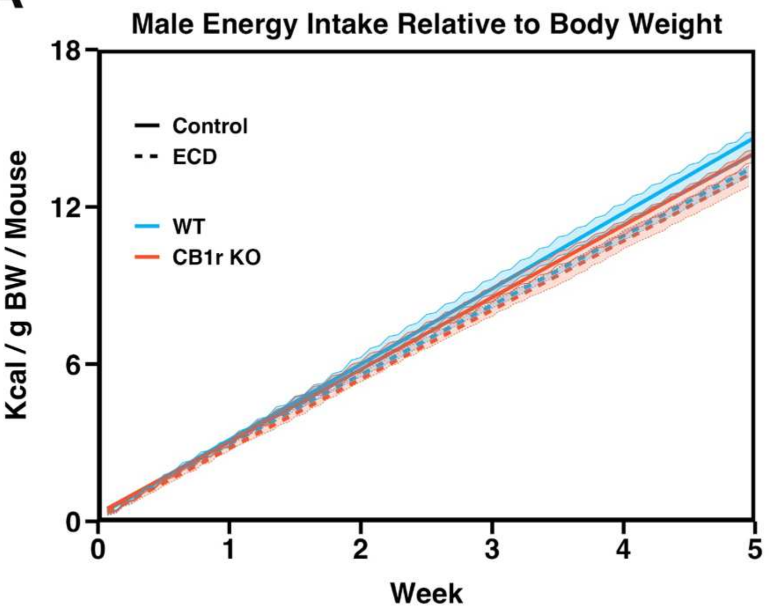

Figure S5

A

Locomotion | Feeding

WT Control

CB1r KO Control

WT ECD

CB1r KO ECD

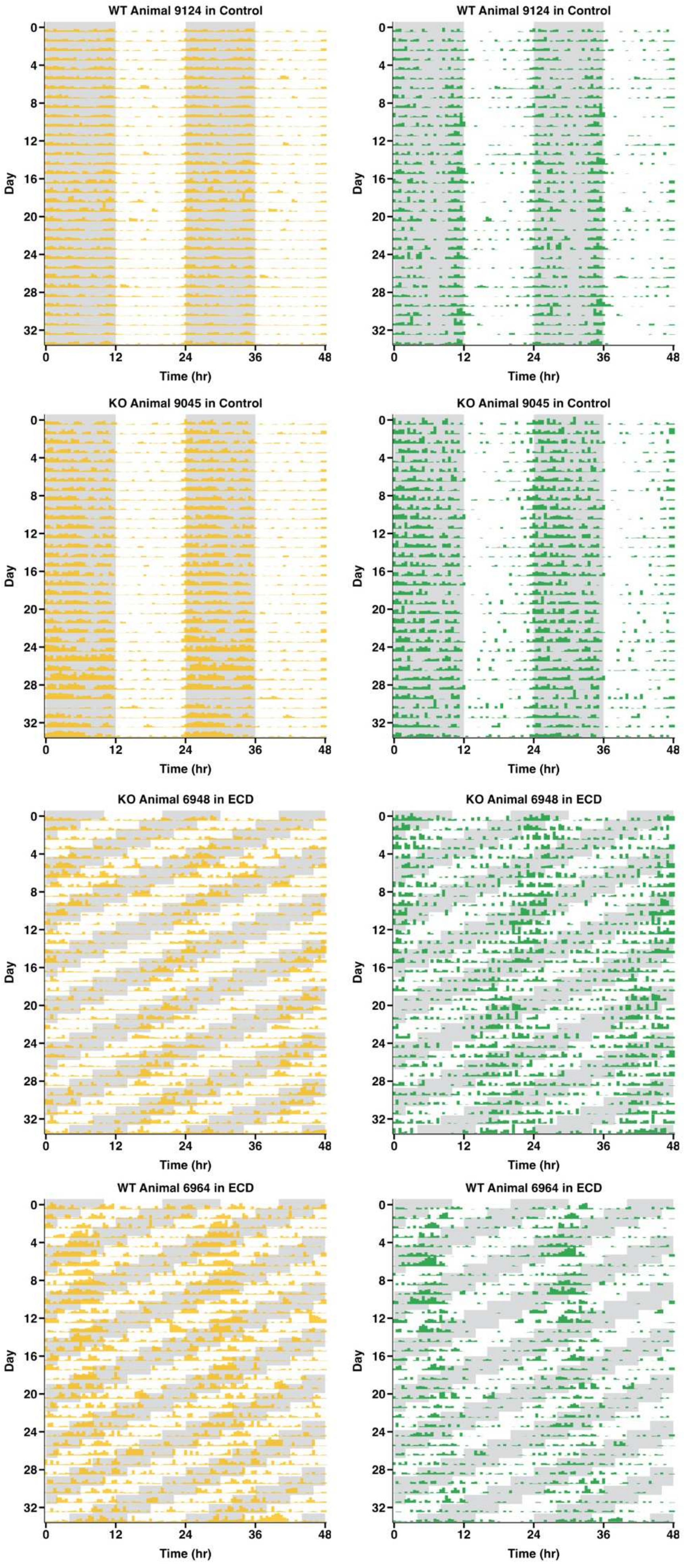

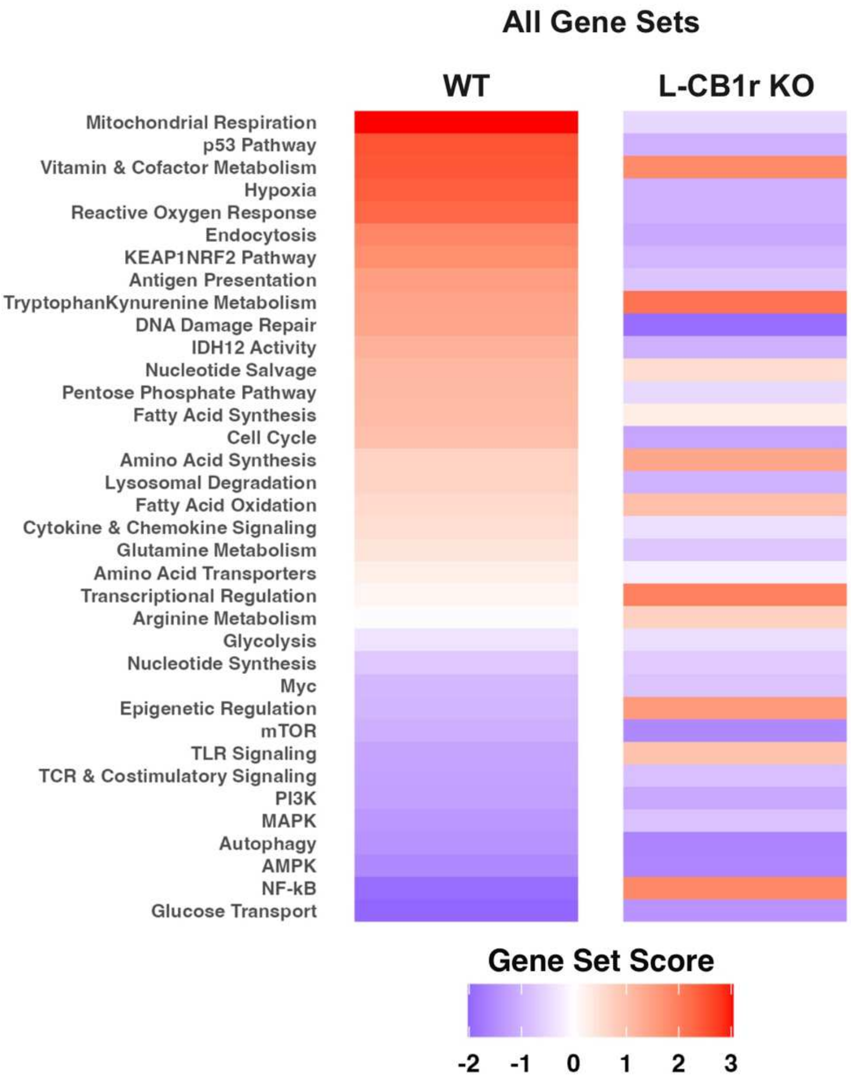
