## Supplemental Figure Captions for "Endocannabinoid signaling is a critical link between circadian desynchronization and metabolic dysfunction"

**Figure S1: ECD has minimal impact on overall levels of circulating or hepatic endocannabinoids, related to Figure 1**

**A – H:** Levels of AEA (left side) and 2-AG (right side) from male (A-D) and female (E-H) plasma and liver samples. Samples were collected at one of the following time points: ZT 0, 4, 8, 12, 16, 20. A 2-way ANOVA was used to test the effect of experimental condition, ZT, and their interaction on endocannabinoid levels. Significant main effects or interaction terms are presented within-figure. Significant follow-up post-hoc tests comparing Control vs ECD, within ZT, are denoted with “*”. *n* = 9-12/experimental condition/ZT.

**I and J:** Scatterplot of AEA and 2-AG levels in plasma (I) and liver (J) from female mice under Control (black circles) and ECD (red triangles) conditions and their linear regressions (lines). Samples were collected at various time points (one of: ZT 0, 4, 8, 12, 16, or 20) and collapsed for correlation analysis. *n* = 30-35/experimental condition/tissue.

**p* < 0.05, ***p* < 0.01, ****p* < 0.001.

**Figure S2: ECD does not induce weight gain in WT or CB1r KO female mice, related to Figure 1**

**A:** Weight gain of female mice recorded manually every 7 days in Control and ECD conditions. Data were analyzed using a linear mixed effects model (*p*_Genotype x Condition x Week_ = 0.786) and within-group follow-up analysis of the effect of ECD performed using estimate of marginal means (*p*_WT_ = 0.155; *p*_KO_ = 0.412). *n* = 57-69/genotype/experimental condition/week.

**Figure S3: ECD alters some, but not all, metabolic related hormones in a CB1r-dependent manner, related to Figures 1 and 6**

**A and B:** Results of the Millipore Mouse Metabolic Hormone Expanded Panel performed on plasma of global CB1r (A) and L-CB1r (B) WT and KO males collected at ZT 12. Main effects or interactions are reported within figure. Significant interaction between genotype and ECD denoted by “#”, significant post-hoc t-test indicated by “*”. *n* = 7-9/genotype/experimental condition.

**C and D:** Triglyceride concentration from a sample of whole liver tissue from global CB1r (C; *p*_Int_ = 0.485, *p*_Geno_ = 0.008) and L-CB1r (D; *p*_Int_ = 0.458, *p*_Geno_ = 0.528) WT and KO males. *n* = 6-9/genotype/experimental condition.

**p* < 0.05, ***p* < 0.01, ****p* < 0.001.

**Figure S4: ECD does not alter energy intake relative to body weight in WT or CB1r KO males, related to Figure 2.**

**A:** Cumulative energy intake, relative to body weight, under Control (solid lines) or ECD (dashed lines) conditions in WT and CB1r KO mice. *p*_Genotype x Condition x Week_ = 0.413; *p*_Genotype x Week_ = 0.094. Data are shown as mean (solid or dashed line) +/- the standard error of the mean (shaded region). For each animal, daily energy intake was standardized to that mouse’s daily body weight, as recorded by the TSE system. Lines indicate group mean, shaded area indicates standard error of the mean.

**Figure S5: Locomotor and feeding rhythms do not align to environmental light cycle in ECD, related to Figure 3.**

**A:** Double plotted actograms depicting locomotion and feeding behavior for representative animals across the entire 35-day experiment. Gray background indicates lights-off, white background indicates lights-on.

**Figure S6: Many gene sets were differentially regulated by ECD in WT and L-CB1r KO mice, related to Figure 7.**

**A:** Heatmap indicating the effect of ECD on gene set score for all gene sets included in the Nanostring Metabolic Pathways Panel. Gene sets were organized in the order of their directed gene score in WTs, from high to low.
